# Differential Behavior of Uncleaved gp160 Human Immunodeficiency Virus Type 1 (HIV-1) Envelope (Env) Glycoproteins

**DOI:** 10.64898/2026.10.14.759227

**Authors:** Stephanie M. Sauve, Mahmoud Moradi

**Affiliations:** Department of Chemistry and Biochemistry, University of Arkansas, Fayetteville, AR 72701

## Abstract

The envelope glycoprotein (Env) of human immunodeficiency virus type 1 (HIV-1) plays important roles in viral entry by coordinating host-cell receptor recognition and membrane fusion. Env is synthesized as a homotrimeric precursor, gp160, which is cleaved into gp120 and gp41 subunits. Although the cleaved form of Env is required for fusion, uncleaved Env remains plentiful on the surface of HIV-1-infected cells. Previous studies have suggested that uncleaved Env is conformationally more flexible than the cleaved form making its dynamics important for understanding the broader structural landscape of HIV-1 Env. In this study, we employed microsecond-long all-atom molecular dynamics (MD) simulations to characterize the dynamics of two structures of uncleaved HIV-1 Env that have been previously assigned to the same conformational state. Our results reveal differences in their conformational behavior, including differences in collective motions, FRET distributions, and trimer symmetry. These findings provide molecular-level insight into the structural plasticity of uncleaved HIV-1 Env and highlight that different structures within the same conformational state can exhibit distinct dynamic behavior.

## Introduction

Human immunodeficiency virus (HIV), the pathogen responsible for acquired immunodeficiency syndrome (AIDS), is a retrovirus that severely damages the immune system. To date, the HIV/AIDS pandemic has claimed the lives of tens of millions of people. ^1–7^ Despite decades of research and the ongoing global health crisis posed by HIV, a cure has yet to be developed. Although antiretroviral therapies can prevent the furthered progression of HIV and transmission of the virus, challenges such as drug resistance, adverse side effects, and an inability to eliminate the virus from the body ^8–15^ are known limitations associated with their use. These limitations highlight the continued need to better understand the molecular mechanisms underlying HIV infection and persistence.

HIV relies on envelope glycoproteins (Env) expressed on the viral surface to recognize host-cell receptors and mediate fusion between the viral and target membranes.^16–23^ Env is initially synthesized in the endoplasmic reticulum as a trimeric gp160 precursor which undergoes proteolytic cleavage by a furin-like protease in the Golgi apparatus. ^24–28^ Once cleaved, the Env protein is a homotrimer of gp120-gp41 heterodimers.^23^ The cleaved Env protein binds the CD4 receptor and a coreceptor, either CCR5 or CXCR4, on target cells to attach to the host cell via its gp120 subunit.^29–35^ Receptor binding causes conformational changes to occur which allow Env to carry out the fusion process via its gp41 subunit.^30–35^ Although cleavage of Env is required for fusion to occur, uncleaved gp160 is also biologically relevant. Uncleaved Env that has bypassed the Golgi is found on the surface of HIV-1-infected cells^27,36,37^ and has been reported to have greater conformational flexibility than cleaved Env.^36,38,39^ This increased flexibility may allow uncleaved Env to access conformations and expose epitopes that are less accessible in the more closed forms of mature Env. ^40^ Thus, uncleaved Env provides an important system for investigating the broader conformational landscape of HIV-1 Env.

Single-molecule Förster resonance energy transfer (smFRET) experiments have shown that HIV-1 Env is conformationally dynamic, sampling multiple conformational states.^23,40–45^ These include a closed pre-triggered state (state-1), a transitional intermediate state (state2), and an open CD4-bound state (state-3).^27,34,39,41,44,46–48^ Among these conformational states, the state-2 intermediate represents a key transitional configuration between the open and closed Env states. More recently, a fourth state called the “occluded intermediate” state has been proposed which occurs between the state-2 and the state-3 conformation.^45^ Env that is unable to be proteolytically processed can access all conformational states, but tends to occupy state-2 and state-3 more often than state-1.^39^ Uncleaved Env therefore provides a useful system for examining the enhanced conformational flexibility of the Env landscape. Characterizing this flexibility at molecular-level resolution is important for determining how structurally distinct conformations within the same broader conformational state can differ in their dynamics and structural organization.

Here, we performed microsecond-long all-atom MD simulations starting from two experimentally characterized uncleaved HIV-1 Env structures,^40^ referred to as S2U1 and S2U2. Although both structures belong to the same broader conformational state, they represent distinct conformations within that ensemble. Through analyses of structural stability, residue-level flexibility, collective motions, computational smFRET, and protomer asymmetry, we characterize differences in their dynamic behavior and conformational sampling. Our findings reveal distinct dynamic behavior for S2U1 and S2U2, providing molecular-level insight into the structural plasticity of uncleaved HIV-1 Env and its conformational heterogeneity.

## Methods

### Molecular Modeling and Simulation Systems

We employed all-atom MD simulations to investigate the conformational dynamics of the uncleaved gp160 precursor HIV-1 Env protein. We designed two gp160 HIV-1 Env simulation systems based on the cryogenic electron microscopy (cryo-EM) structures of the uncleaved HIV-1 JR-FL Env glycoprotein trimer bound to small molecule HIV-1 entry inhibitor BMS-378806 (PDB entry: 7n6u, 4.10 Å resolution, referred to as S2U1 and PDB entry: 7n6w, 4.70

Å resolution, referred to as S2U2).^40^ The small molecule HIV-1 entry inhibitor BMS-378806 and glycans were removed prior to simulation. Representations of the created systems are shown in Figure 1 which was generated using Visual Molecular Dynamics (VMD).^49^

**Fig. 1.**
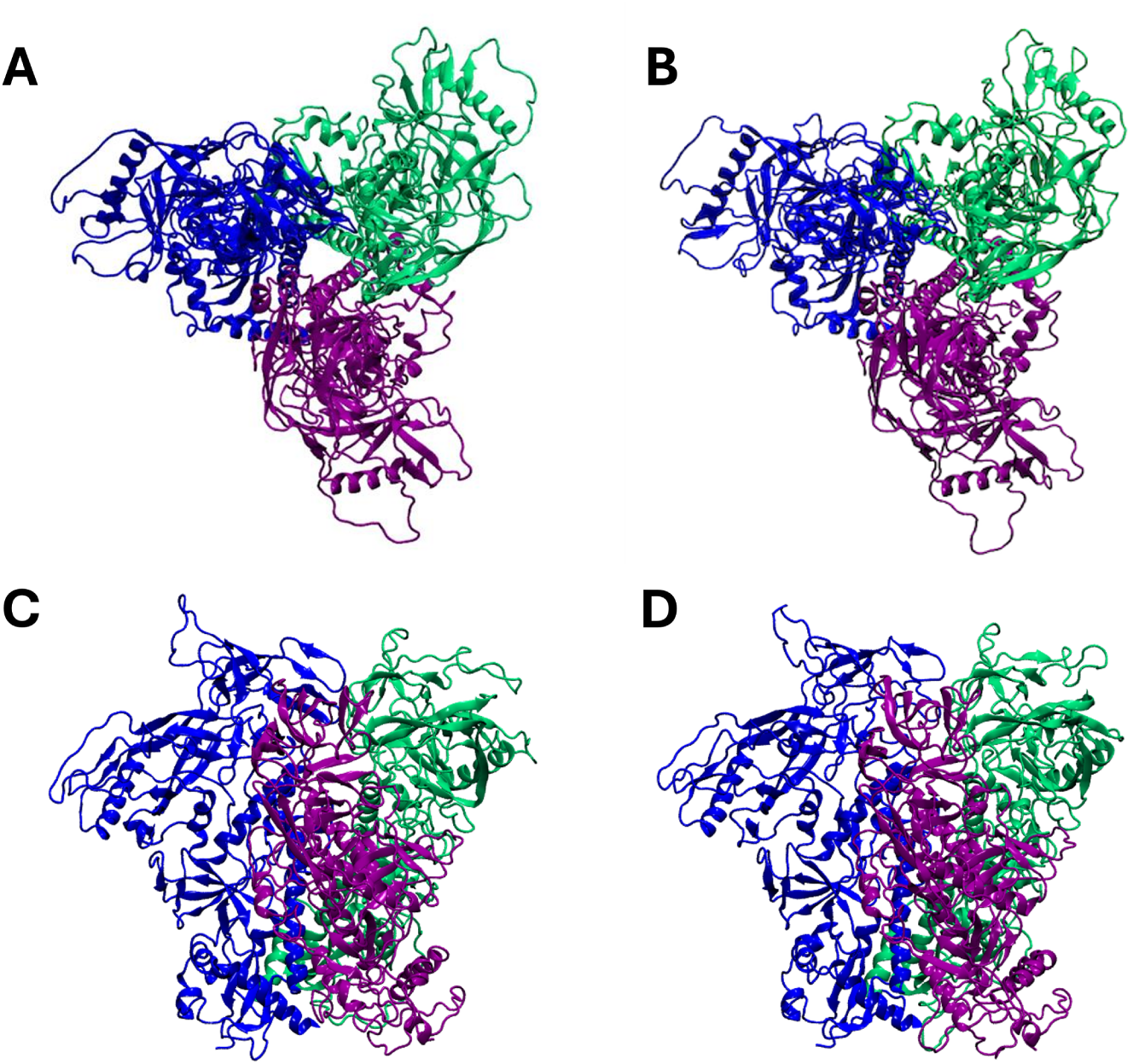
Initial configurations of the uncleaved HIV-1 Env trimer systems used for MD simulations. (A,B) Top views of the S2U1 (left) and S2U2 (right) starting structures, respectively. (C,D) Side views of the S2U1 (left) and S2U2 (right) starting structures, respectively. The three protomers are shown individually with Protomer A colored purple, Protomer B colored blue, and Protomer C colored green.

For both systems, we utilized a Monte Carlo algorithm to model the missing loop regions using the program MODELLER.^50^ The prepared HIV-1 gp160 Env systems were made using the Solution Builder module in CHARMM-GUI^51^ in which 0.15 M NaCl was included (in addition to the counterions used to neutralize the protein) to mimic physiological conditions. The simulation systems were solvated in a cubic box of TIP3P water molecules,^52^ with dimensions of 156 Å *×* 156 Å *×* 156 Å. Each simulation system contained approximately 357,300 atoms. Both S2U1 and S2U2 systems were simulated with ten replicates. All MD simulations were performed in NAMD 2.13, 2.14, and 3.0^53^ using the CHARMM36m force field^54^ for proteins and ions. Energy minimization was initially applied to both of the systems using the conjugate gradient algorithm for 10,000 steps. Production simulations were performed for 2 *µ*s for ten independent replicates of each system and extended to approximately 4.3 *µ*s for the first three replicates of each system, resulting in approximately 54 *µ*s of aggregate simulation data. Due to computational constraints, only three independent replicas per system were extended beyond 2 *µ*s to provide additional conformational sampling.

The equilibration phase was performed in an NVT ensemble, while the production runs were conducted in an NPT ensemble. A time step of 2 fs was utilized, and the temperature was maintained at 310 K using a Langevin thermostat. The Nośe-Hoover Langevin piston method^55^ was used to control pressure at 1 bar.

### Trajectory Analysis

The root-mean-square deviation (RMSD) trajectory tool in VMD^49^ was employed to calculate the RMSD, with C*_α_* atoms considered for these calculations. RMSD distributions were generated from the resulting trajectory values using a bin width of 0.01 Å, followed by kernel density estimation to visualize the conformational ensembles sampled by each system. To assess the flexibility of individual residues, the root-mean-square fluctuation (RMSF) was calculated using C*_α_* atoms, aligning the trajectory against the cryo-EM structure. Average RMSF profiles were calculated for S2U1 and S2U2 ensembles, and differential RMSF values were obtained as:

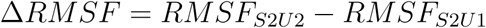

where positive values indicate residues with increased fluctuations in S2U2 relative to S2U1. Principal component analysis (PCA) was performed on the extended system trajectories using the ProDy software,^56^ considering only C*_α_* atoms for the calculations, to characterize the dominant collective motions sampled during the extended simulations.

Computational smFRET analysis was performed by calculating FRET efficiencies from donor-acceptor distances sampled throughout the trajectories. The donor and acceptor labeling positions placed in V1 and V4 were selected based on the experimental smFRET strategy used to characterize Env conformational states.^23,40^ The MD simulations were performed using the native Env sequences without introduction of labeling mutations. For smFRET analysis, separate structural models were generated in which the experimentally labeled residues were mutated to lysine to provide attachment sites for the modeled TCO*^∗^* labeling reagent. These modified structures were aligned to the corresponding simulated Env structures, and the TCO*^∗^* probe was positioned by aligning it to the engineered lysine residues. The donor dye LD555-TTZ and acceptor dye LD655-TTZ were subsequently modeled to the corresponding labeling positions.^23^ Donor-acceptor distances were measured throughout the MD trajectories using the modeled dye positions and were converted to FRET efficiencies according to:

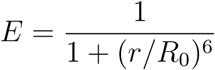

where *r* represents the donor-acceptor distance and *R*_0_ represents the Förster radius of the dye pair.^57–61^ Since Env is a homotrimer, all possible donor-acceptor combinations among the three protomers were considered. At each trajectory frame, the nine possible donor-acceptor pair combinations were converted into individual FRET efficiencies and combined to obtain a weighted FRET efficiency:

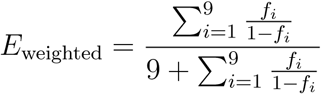

where *f_i_*represents the FRET efficiency calculated for the *i*th donor-acceptor protomer combination. The resulting FRET efficiency values were used to generate distributions representing the conformational ensembles sampled by the S2U1 and S2U2 Env states.

To quantify quaternary rearrangements of the Env trimer, motivated by previously reported differences in protomer organization between the S2U1 and S2U2 conformations,^40^ protomer asymmetry was evaluated by measuring equivalent inter-protomer distances among the three protomers at each trajectory frame. For gp120, distances were measured using the C*α* atoms of residues A333 and Q349. Similarly, the C*α* atoms of residues Y630 and I638 were used for gp41. For both regions, the three inter-protomer distances were calculated in the A-to-C, C-to-B, and B-to-A directions. The asymmetry score was calculated as the sample standard deviation of the three equivalent inter-protomer distances:

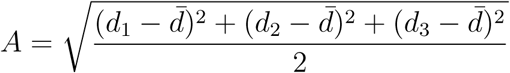

where *d*_1_, *d*_2_, and *d*_3_ represent the three equivalent inter-protomer distances and *d̄* represents their mean value. This metric was used to characterize variations in inter-protomer organization during the simulations. Since the inter-protomer distances were calculated using specific residue pairs, the asymmetry score depends on the direction in which the residue pairs are selected between neighboring protomers. Although the standard deviation is invariant to the ordering of the three measured distances, reversing the direction of the residue-pair selections may produce different asymmetry scores. Therefore, this metric provides a direction-dependent measure of trimer asymmetry rather than a complete description of quaternary organization.

To quantitatively compare the conformational behavior of S2U1 and S2U2, replicate-level statistical analyses were performed on the RMSD, calculated FRET efficiency, and asymmetry data from the ten independent 2 *µ*s simulations of each system. Group-level means and between-replicate standard deviations were subsequently calculated from the ten independent simulations of each system. Differences between S2U1 and S2U2 were evaluated using nonparametric bootstrap resampling. For each comparison, the ten independent replicate-level measurements from each system were sampled independently with replacement over 10,000 bootstrap iterations. The difference between S2U2 and S2U1 was calculated for each bootstrap sample, and the 2.5th and 97.5th percentiles of the resulting distribution were used to estimate 95% confidence intervals. Bootstrap comparisons were performed for the mean RMSD, FRET efficiency, and asymmetry data, the average within-replicate standard deviations, and the between-replicate variability of each observable. All statistical analyses were performed using Python.

## Results and Discussion

### Differential Conformational Dynamics Distinguish S2U1 and S2U2 Env Trimers

To characterize the global conformational behavior of uncleaved HIV-1 Env trimer conformations backbone RMSD values were calculated for each trajectory after alignment to the corresponding starting structure. Ten independent 2 *µ*s long simulations were performed for both S2U1 and S2U2 (Fig. S1), allowing for comparison of the conformational ensembles sampled from each experimentally determined starting conformation.

The RMSD density distributions revealed differences in conformational behavior for the two Env conformations (Fig. 2A–C). The ten S2U1 trajectories exhibited highly overlapping RMSD distributions indicating a consistent conformational response between the independent simulations. A majority of the simulations sampled a relatively narrow range centered around approximately 7 Å (Fig. 2A). In contrast, the S2U2 RMSD distributions displayed greater variability between independent replicates than S2U1. Individual S2U2 replicates sampled broad RMSD ranges spanning approximately 6-10 Å (Fig. 2B). Consequently, the pooled S2U2 RMSD distribution was broader than that of S2U1 and extended toward higher RMSD values indicating greater divergence from the initial conformation (Fig. 2C).

**Fig. 2.**
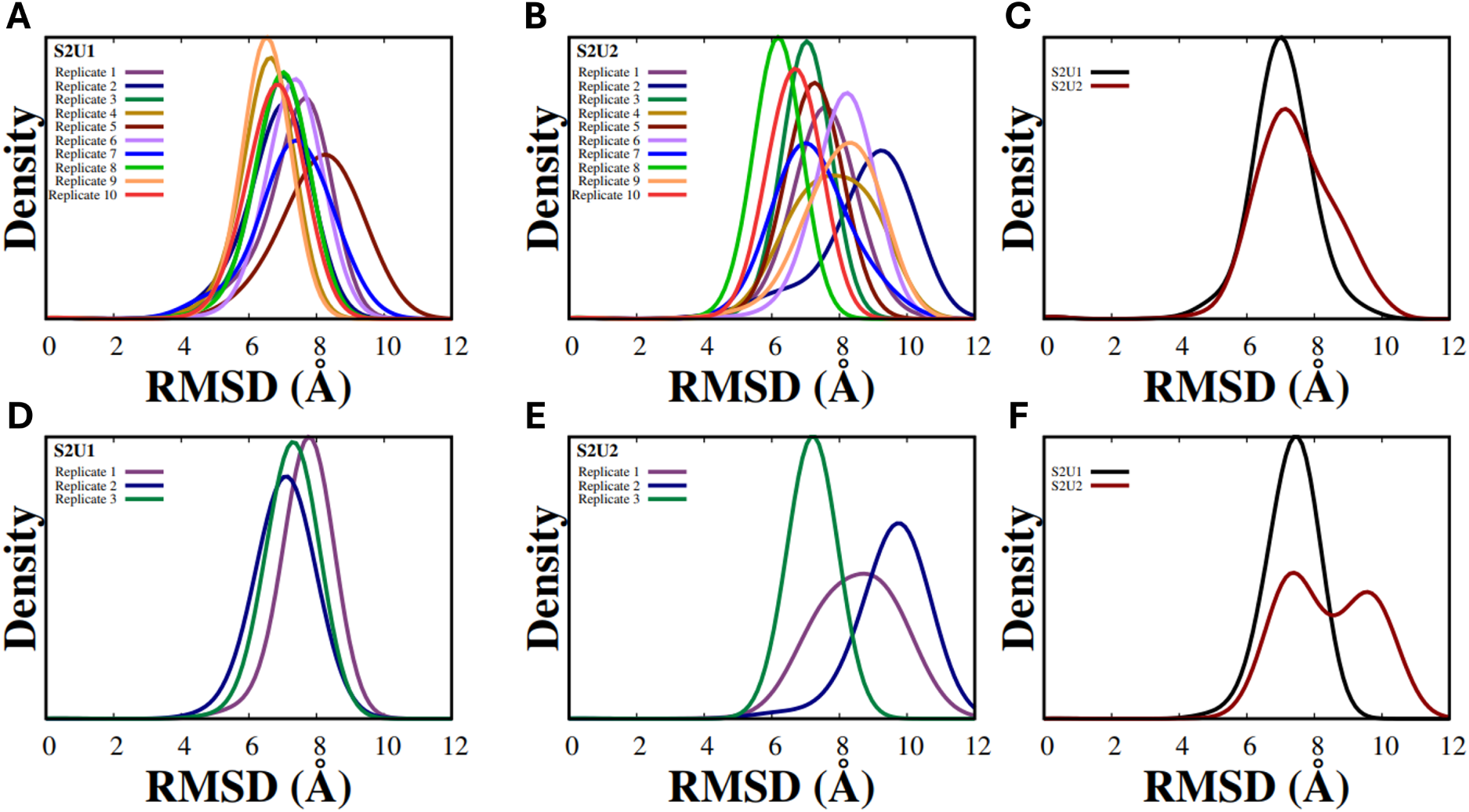
RMSD density distributions calculated for the ten independent S2U1 (A) and S2U2 (B) simulations over the first 2 *µ*s of simulation, with pooled distributions shown in (C). RMSD density distributions for three extended simulations of S2U1 (D) and S2U2 (E) reaching approximately 4.3 *µ*s, with pooled distributions shown in (F). RMSD values were binned using a 0.01 Å interval and smoothed using kernel density estimation.

To quantitatively evaluate these differences, the mean and standard deviation of the RMSD were calculated for each of the ten independent 2 *µ*s production trajectories (Fig. S3). S2U1 exhibited an average RMSD of 7.05 ± 0.48 Å (mean ± between-replicate standard deviation), compared with 7.45 ± 0.77 Å for S2U2. Although the average RMSD was higher for S2U2, bootstrap resampling across independent replicates yielded a mean difference of 0.40 Å (95% confidence interval: -0.15 to 0.93 Å). The average within-replicate RMSD standard deviations were also comparable between S2U1 and S2U2 (0.69 Å and 0.75 Å, respectively), with a difference of 0.06 Å (95% confidence interval: -0.14 to 0.27 Å). Furthermore, S2U2 exhibited greater between-replicate variability in mean RMSD values than S2U1 (0.77 Å versus 0.48 Å), although the bootstrap confidence interval for this difference also included zero (difference: 0.29 Å; 95% confidence interval: -0.10 to 0.64 Å). Together, these findings indicate that while S2U2 exhibits a trend toward greater global structural deviation and variability across independent simulations, these differences were not clearly resolved by the replicate-level statistical comparisons.

To determine whether these differences persisted during longer simulation times, three independent trajectories from each system were extended to approximately 4.3 *µ*s (Fig. S2). These extended trajectories were used to characterize longer-timescale conformational behavior and provide additional observations beyond the ten independent 2 *µ*s simulations used for the quantitative statistical comparisons. The extended simulations further high-lighted differences in the conformational behaviors of the two structures. S2U1 trajectories maintained strongly overlapping RMSD distributions throughout the longer sampling period with little change in the center of the distribution (Fig. 2D). On the other hand, the extended S2U2 simulations diverged into distinct RMSD populations (Fig. 2E). The resulting pooled S2U2 distribution exhibited a broader and multimodal profile compared with the narrower S2U1 pooled distribution (Fig. 2F). Additionally, the S2U2 trajectories underwent larger structural deviations from their respective starting conformations during the extended simulations compared to the S2U1 trajectories.

Together, these results indicate that although both Env structures undergo substantial deviations from their initial conformations, differences in their RMSD distributional behavior emerge across independent simulations. S2U2 exhibits a trend toward greater variability in global structural deviations, particularly during the extended simulations, although the replicate-level statistical comparisons of the 2 *µ*s trajectories did not establish clear differences in average RMSD or variability. These findings suggest that the two uncleaved Env structures may exhibit differences in their conformational dynamics that are not fully captured by average global RMSD measurements alone.

To identify the structural regions contributing to the distinct conformational behavior observed between S2U1 and S2U2, residue-level fluctuations were characterized using RMSF analysis. While both Env conformations exhibited similar overall RMSF profiles, comparison of the ensemble-averaged RMSF values (Fig. S4) revealed localized differences in residue-level dynamics. The difference in RMSF between S2U2 and S2U1 was calculated as ΔRMSF = RMSF_S2U2_ *−* RMSF_S2U1_, where positive values indicate regions with greater fluctuations in S2U2 (Fig. 3). On the other hand, negative values indicate regions with greater fluctuations in S2U1.

**Fig. 3.**
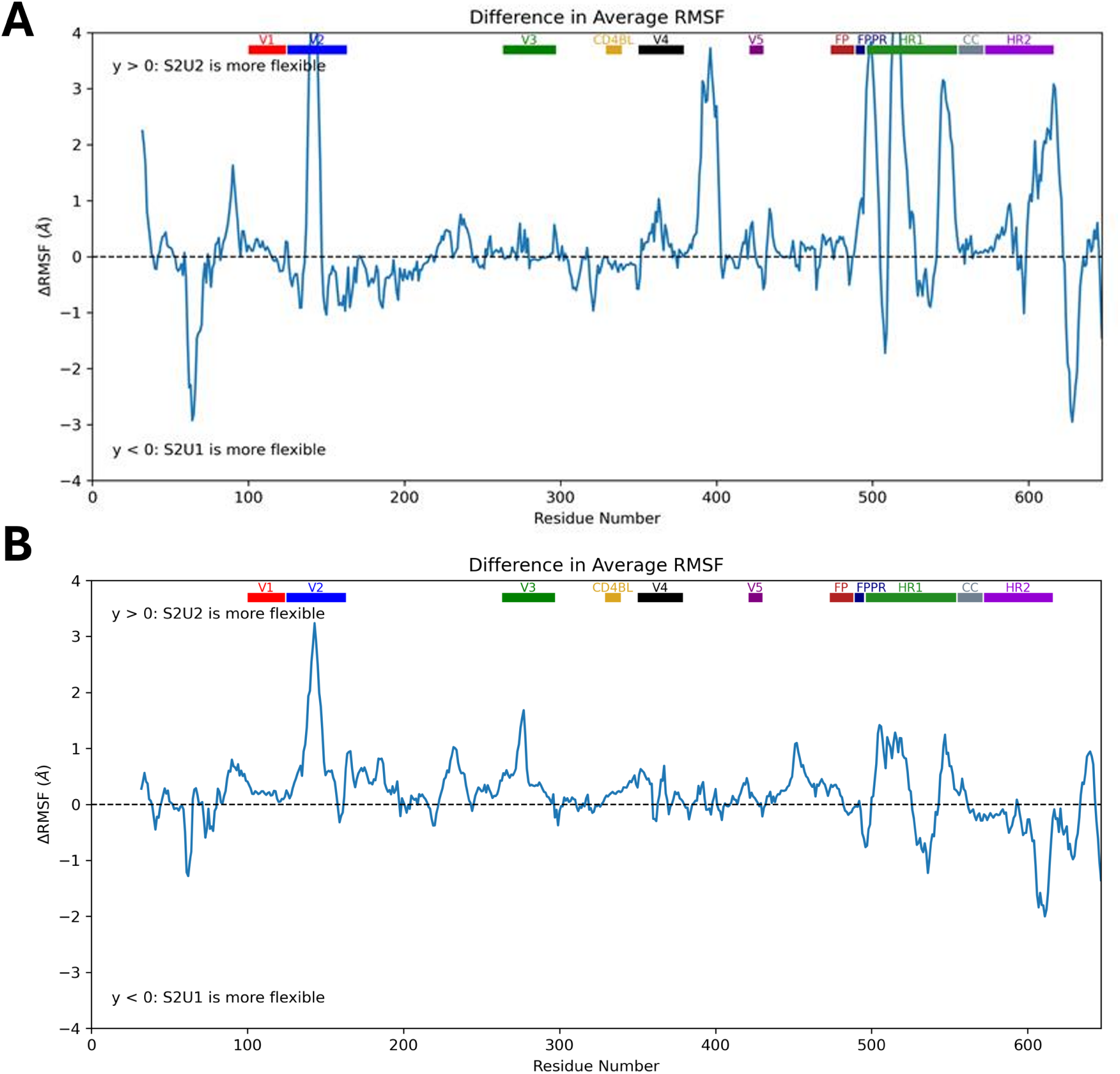
Differential residue-level dynamics of S2U1 and S2U2 Env trimers. The difference in average RMSF between S2U2 and S2U1 was calculated as ΔRMSF = RMSF_S2U2_ *−* RMSF_S2U1_. (A) ΔRMSF profiles obtained from the ten independent 2 *µ*s simulations. (B) ΔRMSF profiles obtained from three extended simulations reaching approximately 4.3 *µ*s. Positive ΔRMSF values indicate residues with greater flexibility in S2U2, whereas negative values indicate residues with greater flexibility in S2U1. Structural regions of the Env trimer are annotated above each profile, including gp120 variable regions (V1, V2, V3, CD4 binding loop (CD4BL), V4, and V5) and gp41-associated regions (fusion peptide (FP), fusion peptide proximal region (FPPR), heptad repeat 1 (HR1), central coiled-coil (CC), and heptad repeat 2 (HR2)).

The ΔRMSF profiles revealed that the enhanced conformational variability of S2U2 was concentrated within specific structural regions of Env. In S2U2, increased gp120 fluctuations were observed within the V1/V2 region. Our findings are consistent with the greater conformational variation in S2U2 reported for the trimer association domain which includes the V1/V2 region in the original structural study.^40^ It is hypothesized that greater structural variation in this region is what leads to a greater extent of asymmetry in S2U2 than S2U1.^40^ Furthermore, differences were observed within the gp41-associated regions, including the fusion peptide proximal region (FPPR) and the heptad repeat region 1 (HR1). These findings are also consistent with the greater structural variation among the gp41 protomers of S2U2 compared with S2U1 reported in the original structural study.^40^ Overall, these regions are involved in regulating Env’s conformations, suggesting that localized changes in residue-level dynamics contribute to the altered conformational sampling observed for S2U2.

Analysis of the extended trajectories revealed that some regional differences in residue-level flexibility persisted during longer simulations (Fig. 3B). However, differences in the gp41 region are much less pronounced in the extended simulations than in the initial simulations, indicating that some of the initially observed gp41 differences were not maintained to the same extent over longer timescales. Nevertheless, the extended simulations continued to show that the two structures displayed different patterns of regional fluctuations. Together with the RMSD distributions, these results suggest that S2U2 accesses a broader conformational landscape with differences in the dynamics of particular structural regions.

### Distinct FRET Sampling Behaviors of S2U1 and S2U2 Within the Same State-2 Conformational Ensemble

It is known that Env proteins predominantly sample state-2 conformations and that breaking symmetry within the Env trimer has been associated with adoption of the state-2 conformational state.^44^ Therefore, both S2U1 and S2U2 were characterized as distinct conformations within the state-2 conformational state in the study from which these structures were obtained.^40^ To determine whether the differences in conformational behavior observed from RMSD and RMSF analyses were reflected in FRET efficiencies, computational smFRET analysis was performed on the MD trajectories. FRET efficiencies were calculated from the donor-acceptor distances sampled throughout each simulation and used to characterize the distance-dependent conformational behavior of the S2U1 and S2U2 Env trimers (Fig. S5).

The calculated FRET distributions revealed differences in the conformational preferences of the two Env structures (Fig. 4). Across the ten independent 2 *µ*s simulations, S2U1 exhibited broad FRET distributions, with individual replicas sampling a wide range of FRET efficiencies. In contrast, S2U2 displayed higher FRET states across independent simulations, resulting in a collective distribution shifted toward higher FRET efficiencies compared to S2U1. Importantly, the FRET behavior observed for both structures were consistent with their previous assignment to the state-2 conformational state. At the same time, the distinct FRET distributions indicate that S2U1 and S2U2 populate different ensembles of donor-acceptor distance states within this conformational state.

**Fig. 4.**
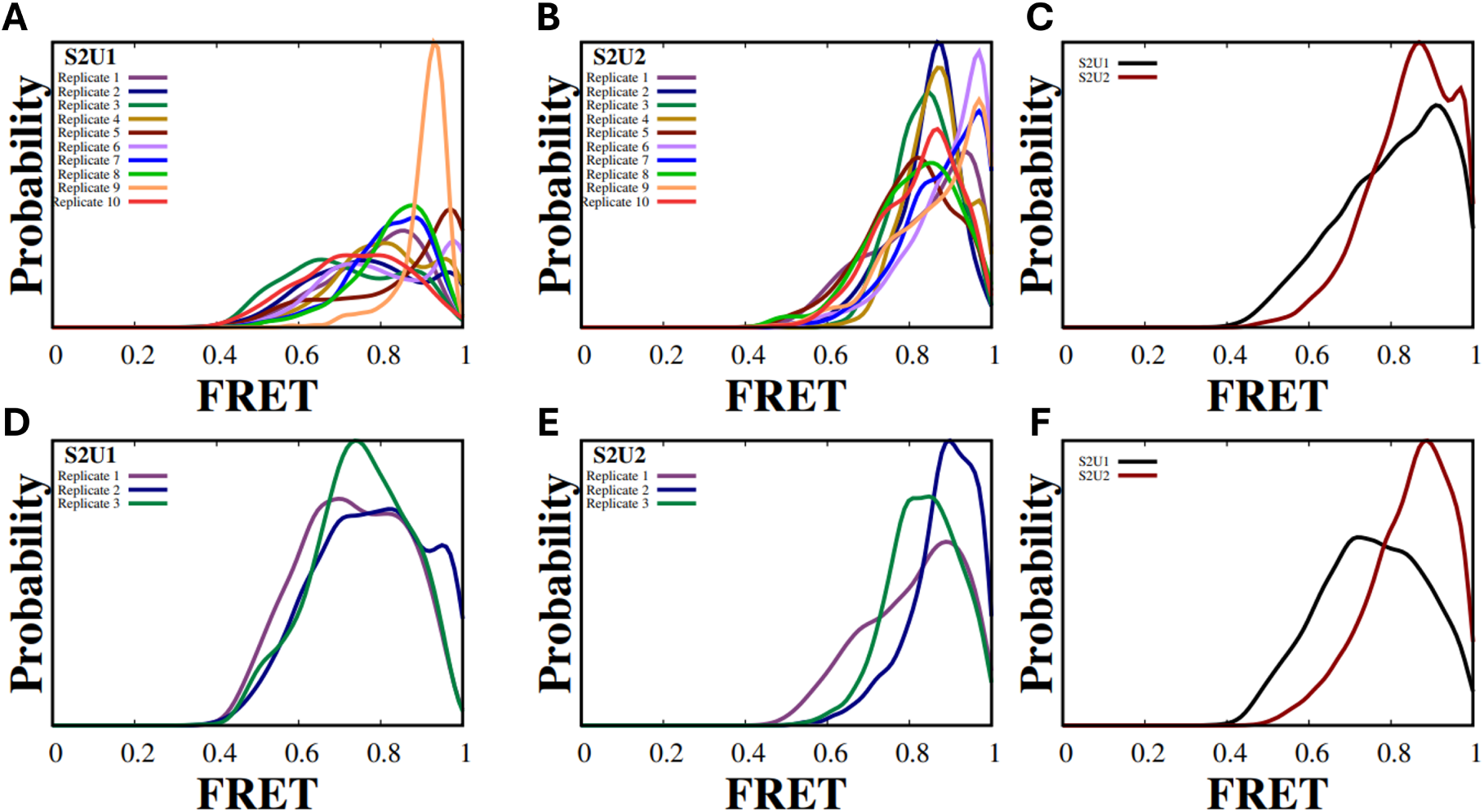
Computational smFRET efficiency distributions reveal differences in donor-acceptor distance sampling between S2U1 and S2U2. (A-C) FRET distributions obtained from ten independent 2 *µ*s simulations of S2U1 and S2U2. (A) Individual replicate FRET distributions for S2U1, (B) individual replicate FRET distributions for S2U2, and (C) combined ensemble distributions comparing S2U1 and S2U2. (D-F) FRET distributions obtained from three extended simulations reaching approximately 4.3 *µ*s. (D) Individual replicate FRET distributions for extended S2U1 simulations, (E) individual replicate FRET distributions for extended S2U2 simulations, and (F) combined ensemble distributions.

Although the FRET distributions differed between the two structures, both S2U1 and S2U2 predominantly sampled high FRET efficiencies, particularly within the range of approximately 0.8–1.0. Since the relationship between donor–acceptor distance and FRET efficiency is nonlinear, efficiencies approaching 1.0 are less sensitive to changes in distance. Consequently, the high-efficiency region compresses the underlying distance distributions, potentially obscuring differences in conformational sampling between the two structures.

To quantitatively evaluate these differences, the mean and standard deviation of the calculated FRET efficiencies were determined for each of the ten independent 2 *µ*s production trajectories (Fig. S6). S2U1 exhibited an average FRET efficiency of 0.805 ± 0.055 (mean ± between-replicate standard deviation), compared with 0.846 ± 0.031 for S2U2. Bootstrap resampling across independent replicates yielded a mean difference of 0.041 (95% confidence interval: 0.004–0.078), supporting higher average FRET efficiencies in S2U2. In contrast, S2U1 exhibited greater within-replicate variation in FRET efficiency, with an average standard deviation of 0.117 compared with 0.096 for S2U2 (difference: -0.020; 95% confidence interval: -0.038 to -0.001). S2U1 also displayed greater between-replicate variability in mean FRET efficiency (0.055 versus 0.031), although the bootstrap confidence interval for this difference included zero. These quantitative findings indicate that S2U1 samples a broader range of calculated FRET efficiencies within individual trajectories, while S2U2 preferentially samples higher average FRET efficiencies.

To evaluate whether these differences were maintained with increased sampling, the computational smFRET analysis was repeated for the three extended simulations reaching approximately 4.3 *µ*s (Fig. 4D-F). The extended trajectories preserved the same trend observed in the initial simulations, with S2U1 maintaining broader FRET sampling while S2U2 remained populated at higher FRET efficiencies. These longer simulations provide additional observations supporting differences in the FRET sampling behavior of the two structures.

Together with the RMSD and RMSF analyses, these results demonstrate that S2U1 and S2U2 exhibit differences in their conformational dynamics despite being characterized as distinct structures within the same state-2 ensemble. While the RMSD analysis did not establish clear differences in average global structural deviations, the computational FRET analysis revealed differences in both the average FRET efficiencies and their fluctuations across independent simulations. These findings suggest that conformations assigned to the same experimentally characterized smFRET state can exhibit distinguishable distance-dependent dynamics that are not fully captured by global structural measurements alone.

### S2U1 and S2U2 Access Distinct Collective Motions

To further characterize the global motions underlying the distinct conformational ensembles observed for S2U1 and S2U2, PCA was performed on the extended MD trajectories. The first three principal components accounted for approximately 52% of the total variance, with PC1, PC2, and PC3 contributing 26%, 14%, and 12%, respectively (Fig. S7A). Projection of the trajectories onto the first two principal components revealed distinct patterns of conformational sampling between the two Env variants (Fig. 5). Additional projections involving PC3 are shown in Figure S7.

**Fig. 5.**
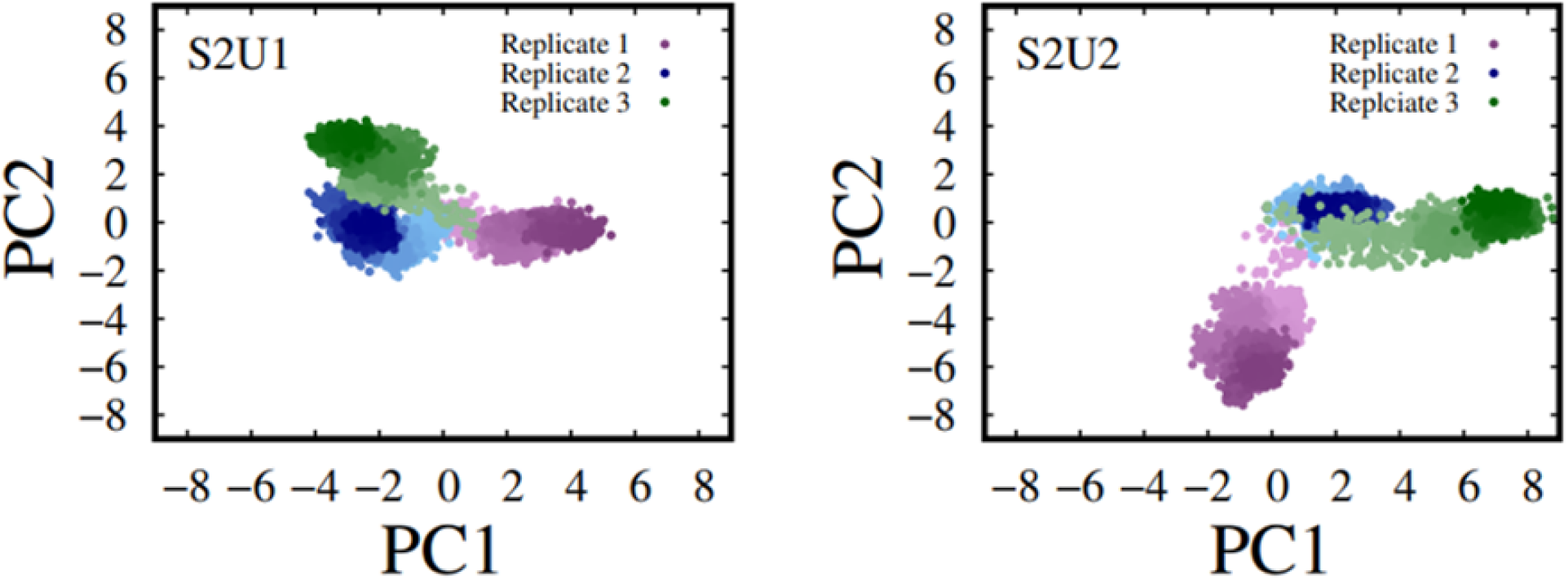
PCA reveals distinct collective conformational sampling of S2U1 and S2U2 Env trimers. PCA was performed on the extended MD trajectories to characterize the dominant collective motions sampled by each Env variant. The projections of the trajectories onto the first two principal components are shown for S2U1 (left) and S2U2 (right). PC1 and PC2 account for 26% and 14% of the total variance, respectively (40% cumulative variance among the first two principal components). Each color represents an independent simulation replica.

The S2U1 trajectories occupied relatively compact regions within the principal component space, indicating that the trimer sampled a more restricted subset of conformational states during the extended simulations. In contrast, S2U2 trajectories explored a broader region of the conformational landscape, with individual replicas accessing distinct regions along PC1 and PC2. The broader sampling observed between S2U2 trajectories suggests enhanced heterogeneity in the collective motions sampled by the trimer.

The differences in conformational sampling observed in the PCA projections are consistent with the variations in RMSD distributions and localized residue-level dynamics identified through RMSF analysis. Together, these analyses suggest that S2U1 and S2U2 exhibit differences in their collective conformational behavior during the extended simulations.

### Protomer Asymmetry Reveals Variability in Env Trimer Organization

To further characterize differences in the trimer organization between S2U1 and S2U2, we quantified protomer asymmetry by measuring the variability of equivalent inter-protomer distances across the three protomers following the approach done by Zhang et al. ^40^ This analysis provides a direct measure of deviations from a symmetric trimer arrangement and allows assessment of how each Env structure samples quaternary conformations.

Representative trimer geometries used for the asymmetry analysis are shown in Figure 6(A,B). For each trajectory frame, the standard deviation of the three equivalent interprotomer distances was calculated as the protomer asymmetry score. To quantitatively compare the two structures, the mean and standard deviation of the gp120 and gp41 asymmetry scores were determined for each of the ten independent 2 *µ*s simulations (Fig. S9). For gp120, S2U1 exhibited an average asymmetry score of 7.36 ± 2.76 Å (mean ± between-replicate standard deviation), compared with 7.86 ± 3.10 Å for S2U2. Although S2U2 exhibited a slightly higher average gp120 asymmetry score, bootstrap resampling across independent replicates yielded a mean difference of 0.50 Å (95% confidence interval: -1.91 to 2.96 Å). Similarly, the average gp41 asymmetry scores were comparable between S2U1 and S2U2 (4.74 ± 3.01 Å and 4.80 ± 2.22 Å, respectively), with a mean difference of 0.06 Å (95% confidence interval: -2.15 to 2.25 Å). The within-replicate fluctuations in both gp120 and gp41 asymmetry were also comparable between the two structures, with bootstrap confidence intervals that included zero. These findings indicate that the initial 2 *µ*s simulations did not establish clear differences in average protomer asymmetry between S2U1 and S2U2.

**Fig. 6.**
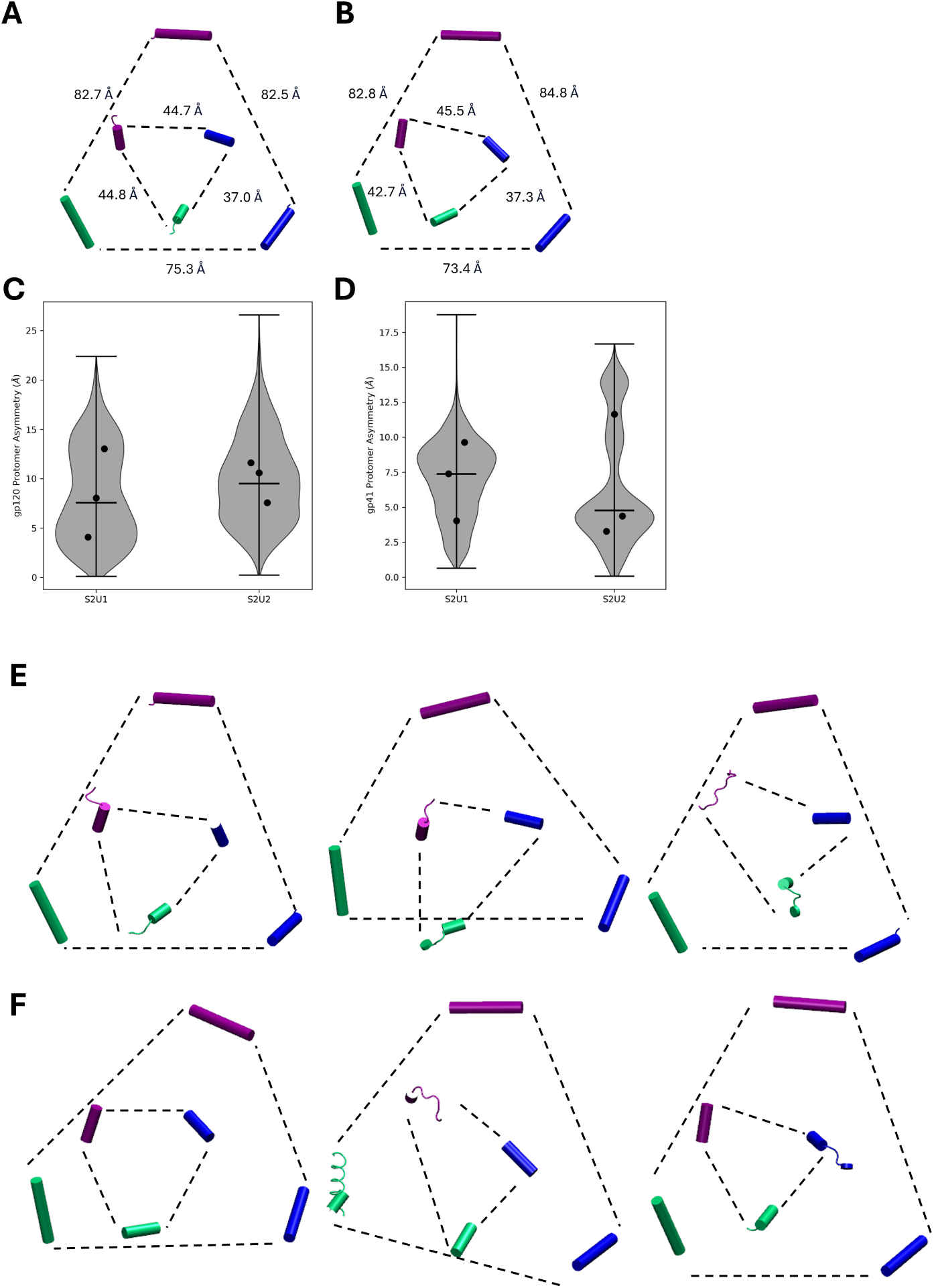
Protomer asymmetry analysis of the S2U1 and S2U2 uncleaved HIV-1 Env trimers. (A,B) Representative trimer geometries of S2U1 and S2U2 showing the interprotomer distances used to quantify protomer asymmetry. Protomer A, B, and C are colored purple, blue, and green, respectively. (C,D) Violin plots showing the distributions of gp120 (C) and gp41 (D) protomer asymmetry values calculated from the extended MD simulations. The violin distributions represent the combined data from three independent replicas for each system, with black dots indicating the mean asymmetry values of individual replicas and horizontal lines indicating the median values.(E,F) Representative endpoint conformations from three independent extended simulations of S2U1 (E) and S2U2 (F), illustrating the range of trimer geometries sampled by each system. Protomer colors are maintained as in panels A and B.

To examine the asymmetry behavior over longer timescales, the analysis was extended to the three independent simulations of each structure reaching approximately 4.3 *µ*s (Fig. S8). The extended trajectories revealed differences in the asymmetry distributions sampled by the two structures. The gp120 asymmetry distribution was shifted toward larger values in S2U2, with the three extended replicates exhibiting higher average asymmetry scores than their S2U1 counterparts (Fig. 6C). This suggests that the extended S2U2 trajectories sampled more asymmetric gp120 arrangements within the analyzed subset.

In contrast, gp41 displayed a broader and more heterogeneous asymmetry distribution in S2U2. Although the dominant population occurred at lower asymmetry values compared with S2U1, S2U2 sampled an additional high-asymmetry population that was absent in S2U1 (Fig. 6D). This suggests that S2U2 does not uniformly exhibit greater gp41 asymmetry, but instead samples a wider range of gp41 protomer arrangements in these extended simulations.

Consistent with the extended-simulation asymmetry distributions, representative endpoint conformations illustrated differences in the trimer geometries sampled by S2U1 and S2U2 (Fig. 6E,F). Although the quantitative comparisons across ten independent 2 *µ*s simulations did not establish clear differences in average protomer asymmetry, the extended trajectories revealed differences in the distributions of quaternary arrangements sampled by the two structures. Together with the RMSD, RMSF, computational smFRET, and PCA analyses, these findings suggest that S2U1 and S2U2 exhibit differences in specific aspects of their conformational dynamics, highlighting how two structures within the same experimentally characterized state-2 ensemble can display distinct patterns of conformational sampling.

Several limitations should be considered when interpreting the findings of this study. The simulations were initiated from cryo-EM structures resolved at 4.1 and 4.7 Å for S2U1 and S2U2, respectively, with missing loop regions reconstructed using MODELLER. Consequently, uncertainties in the starting structures and modeled regions may influence the conformational dynamics observed during the simulations. Additionally, glycans were not included, the BMS-378806 inhibitor present in the experimental structures was removed, and the simulations were performed without an explicit lipid membrane due to only containing part of the gp41 domain. These differences from the experimental conditions may influence Env conformational behavior. Furthermore, although the simulations extended to microsecond timescales, they remain substantially shorter than the timescales accessible through experimental smFRET measurements, limiting the extent of conformational sampling. This sampling limitation is also reflected in the gp41 RMSF analysis, where differences between S2U1 and S2U2 observed during the initial 2 *µ*s simulations became less pronounced in the extended trajectories. Therefore, some of the observed conformational differences may be sensitive to simulation length.

## Conclusion

In summary, we used extensive all-atom MD simulations to characterize the conformational dynamics of two experimentally determined uncleaved HIV-1 Env structures, S2U1 and S2U2.^40^ Across multiple independent simulations and complementary analyses, we demonstrate that these two structures display differences in their conformational dynamics despite being characterized as conformations within the same state-2 ensemble. Overall, our findings highlight that different structures within the same broader conformational state can exhibit different dynamic behavior.

RMSD and RMSF analyses revealed differences in the structural fluctuations and conformational behavior of S2U1 and S2U2, although quantitative RMSD comparisons did not establish clear differences in their average global structural deviations. Computational FRET analysis further revealed differences in their donor-acceptor distance-dependent behavior, with S2U2 exhibiting higher average calculated FRET efficiencies and S2U1 displaying greater fluctuations in FRET efficiency. PCA of extended simulations showed that S2U2 explored a broader conformational space compared with S2U1. Furthermore, analysis of inter-protomer organization revealed greater heterogeneity in the quaternary arrangements sampled by S2U2 during the extended simulations.

These findings provide a dynamic perspective on the structural differences observed between the S2U1 and S2U2 cryo-EM structures and suggest that subtle differences in experimentally resolved conformations correspond to distinct underlying conformational sampling. Together, this work highlights the importance of considering the dynamic behavior of Env trimers beyond their static structural representations and provides molecular-level insight into the conformational variability of uncleaved HIV-1 Env.

## Supporting Information Available

Figures and information in the Supplemental Information provide additional analysis based on our MD simulations not discussed in the manuscript.

## Acknowledgement

The research reported in this work was supported by the National Institute of General Medical Sciences of the National Institutes of Health under award number R35GM147423, as well as by the National Science Foundation grant CHE 1945465 and the Arkansas Biosciences Institute. This research is part of the Frontera computing project by LRAC at the Texas Advanced Computing Center (TACC) through Grant CHE21003, made possible by the National Science Foundation award OAC-1818253. Anton 3 computer time was provided by the Pittsburgh Supercomputing Center (PSC) through Grant 1R24GM154042 from the National Institutes of Health. The Anton 3 machine at PSC is made available by D. E. Shaw Research. This work is also supported by the Arkansas High-Performance Computing Center which is funded through multiple National Science Foundation grants and the Arkansas Economic Development Commission.

