## Supplemental Information for "Differential Behavior of Uncleaved gp160 Human Immunodeficiency Virus Type 1 (HIV-1) Envelope (Env) Glycoproteins"

To evaluate whether the conformational behavior observed during the initial 2 $\mu$s simulations (Fig. S1) was maintained at longer timescales, three independent replicas of both S2U1 and S2U2 were extended to approximately 4.3 $\mu$s. RMSD trajectories for the extended simulations are shown in Figure S2. All extended trajectories exhibited an initial increase in RMSD followed by fluctuations around relatively stable values, consistent with the behavior observed during the 2 $\mu$s simulations.

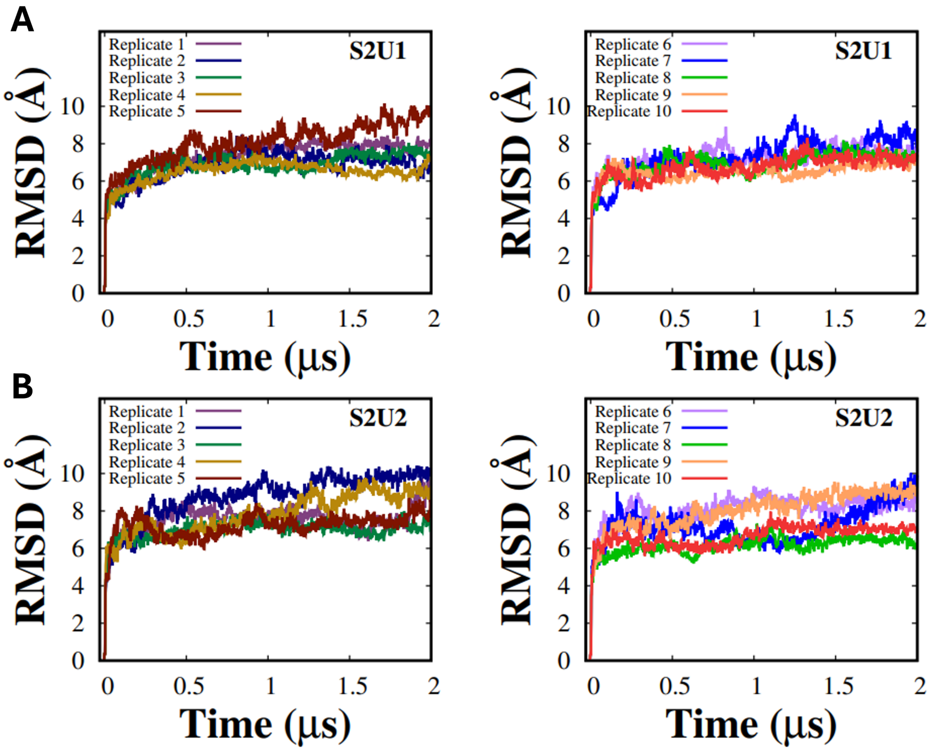


**Fig. S1.** RMSD time series plots for the 2 $\mu$s MD simulations of the uncleaved HIV-1 Env trimer. (A) S2U1 and (B) S2U2, with trajectories shown for all ten independent replicas. For clarity, replicas 1-5 (left) and 6-10 (right) are displayed in separate panels within each system.

The extended S2U1 replicas remained within a similar RMSD range throughout the additional simulation time (Fig. S2A), whereas the S2U2 replicas displayed greater variability between independent trajectories (Fig. S2B). These extended simulations preserve the differences observed between the two Env states during the shorter simulations and provide additional sampling of the conformational ensembles used for subsequent analyses.

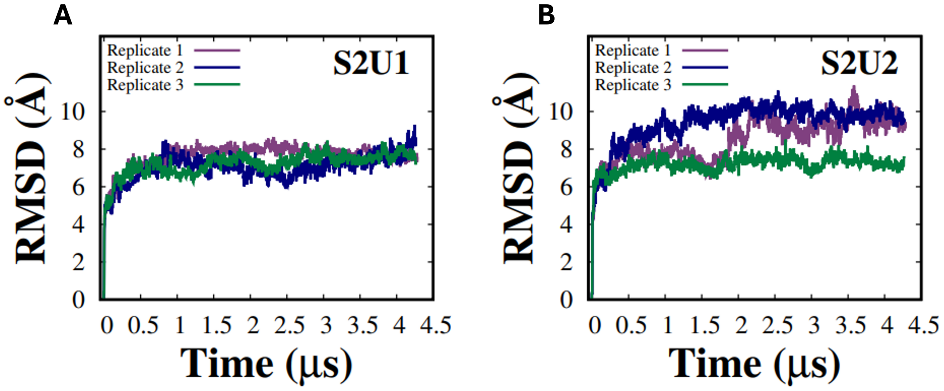


**Fig. S2.** RMSD time series plots for the extended MD simulations of the uncleaved HIV-1 Env trimer, showing three independent replicas for each system extended to approximately 4.3 $\mu$s. S2U1 in shown in panel A and S2U2 is shown in panel B.

To further characterize the global structural deviations of S2U1 and S2U2, the mean backbone RMSD was calculated for each of the ten independent 2 $\mu$s production simulations (Fig. S3). S2U1 exhibited an average RMSD of 7.05 ± 0.48 Å (mean ± between-replicate standard deviation), compared with 7.45 ± 0.77 Å for S2U2. Although S2U2 exhibited a slightly higher average RMSD and greater variability among independent simulations, the 95% bootstrap confidence interval for the difference in mean RMSD included zero (-0.15 to 0.93 Å). These results indicate that the average magnitudes of global structural deviation were broadly comparable between the two systems.

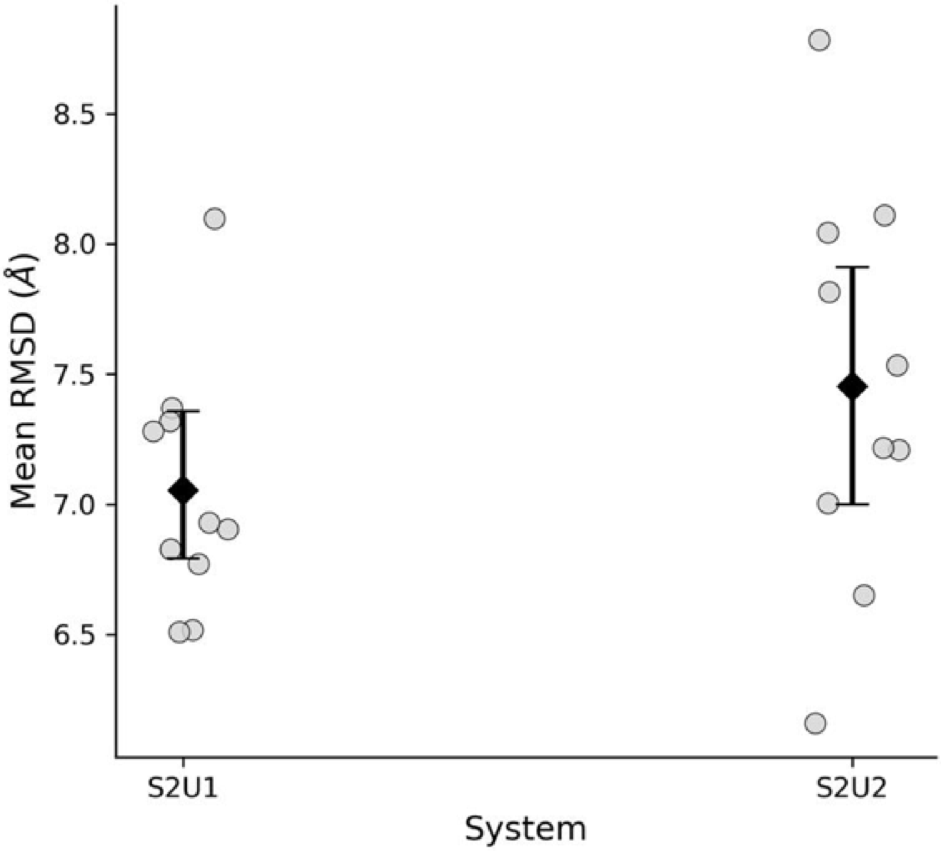


**Fig. S3.** Comparison of mean RMSD values across independent S2U1 and S2U2 simulations. Each grey point represents the mean backbone RMSD calculated from the production frames of an individual 2 $\mu$s simulation. Black diamonds represent the means across ten independent simulations, with error bars indicating 95% bootstrap confidence intervals obtained by resampling replicate means (10,000 iterations).

To assess whether the longer simulation times altered the observed residue-level dynamics, average RMSF profiles were calculated from both the initial 2 $\mu$s simulations and the extended 4.3 $\mu$s simulations for S2U1 and S2U2 (Fig. S4). The RMSF profiles obtained from the extended trajectories (Fig. S4C-D) showed similar overall fluctuation patterns compared with the corresponding 2 $\mu$s simulations for both Env states (Fig. S4A-B).

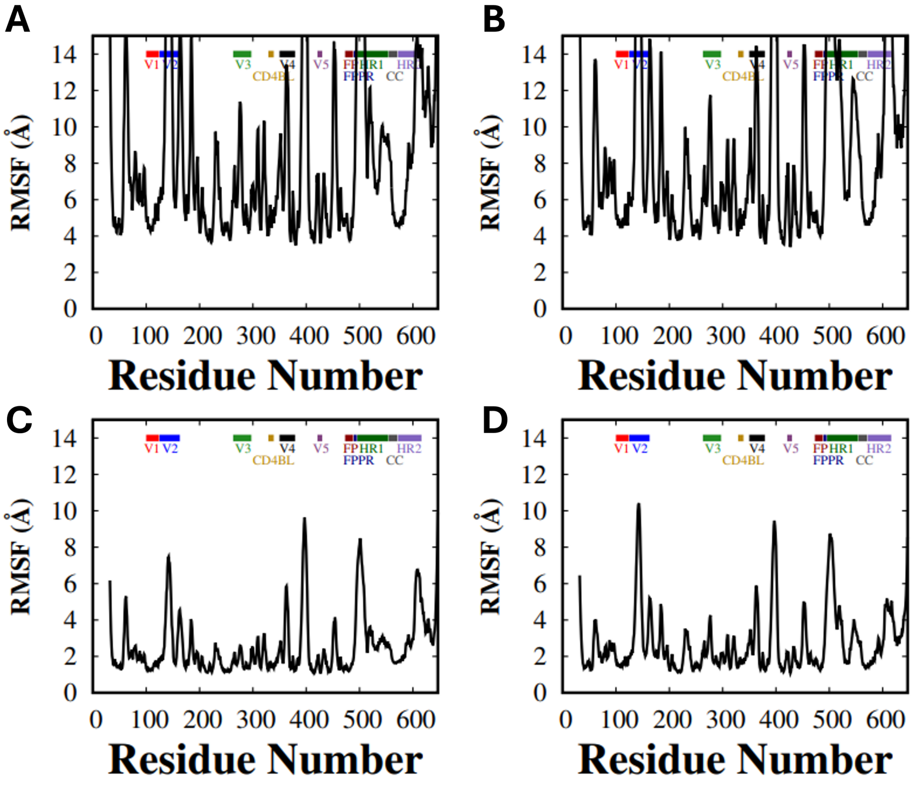


**Fig. S4.** Average RMSF profiles for the uncleaved HIV-1 Env trimer simulations. (A) S2U1 and (B) S2U2 RMSF profiles calculated from the 2 $\mu$s simulations.(C) S2U1 and (D) S2U2 RMSF profiles calculated from the extended 4.3 $\mu$s simulations.

In both S2U1 and S2U2, elevated fluctuations were primarily localized to flexible loop regions, while the overall residue-level fluctuation profiles remained consistent between simulation lengths.

To evaluate the behavior of the computational smFRET signals over time, FRET efficiencies were monitored throughout each MD trajectory (Fig. S5). The 2 $\mu$s simulations for both S2U1 and S2U2 exhibited continuous fluctuations between different FRET efficiency values, reflecting the dynamic nature of the Env trimer conformational ensembles (Fig. S5A-D).

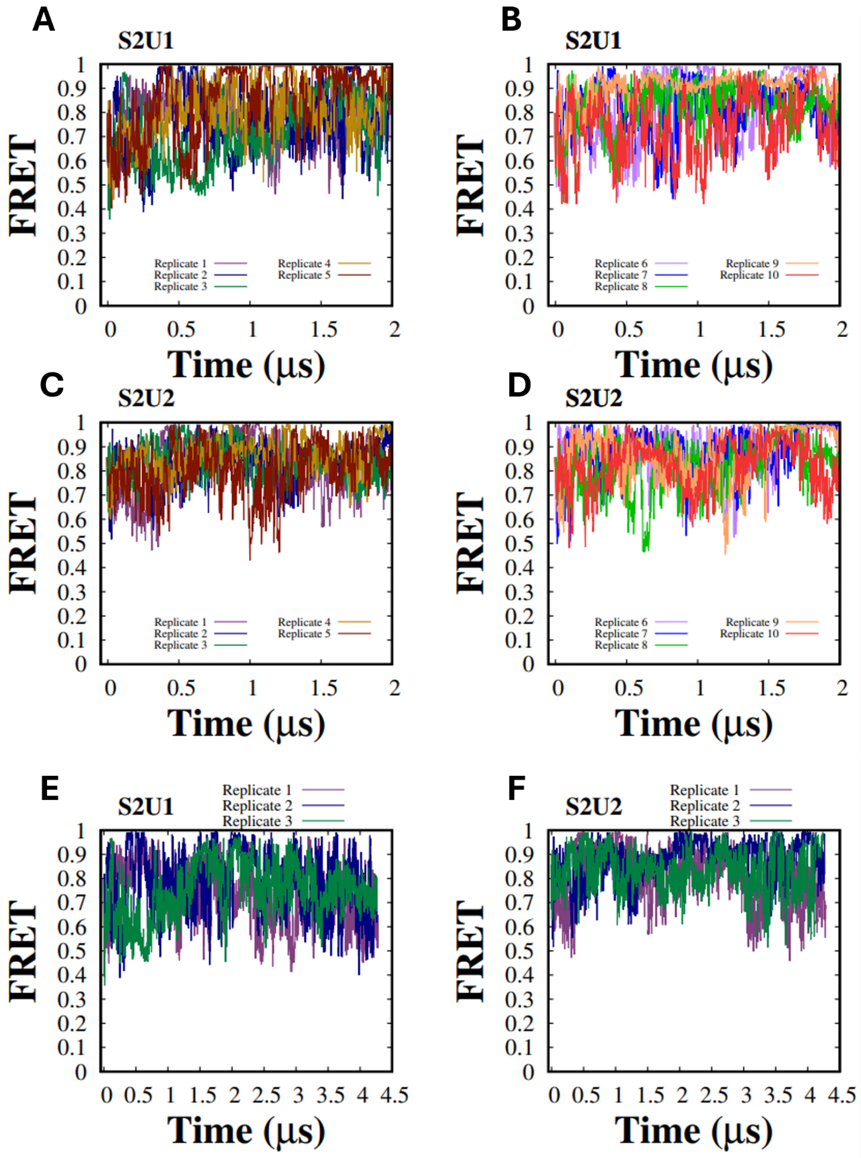


**Fig. S5.** Time evolution of computational smFRET efficiencies for the uncleaved HIV-1 Env trimer simulations. (A,B) smFRET trajectories for the ten independent 2 $\mu$s simulations of S2U1. (C,D) smFRET trajectories for the ten independent 2 $\mu$s simulations of S2U2. (E,F) smFRET trajectories for the three extended $\sim$4.3 $\mu$s simulations of S2U1 (left) and S2U2 (right), respectively. Each trace represents the calculated FRET efficiency sampled throughout the MD trajectory. The trajectories demonstrate the conformational fluctuations underlying the smFRET distributions presented in the main text.

The extended simulations maintained similar FRET sampling behavior observed during the initial simulations, with S2U1 and S2U2 exploring distinct ranges of FRET efficiencies (Fig. S5E-F). These trajectory-level observations support the FRET distributions presented in the main text and demonstrate that the calculated FRET profiles arise from sustained sampling of the underlying conformational ensembles rather than isolated structural events.

To further characterize differences in the FRET sampling behavior of S2U1 and S2U2, the mean and standard deviation of the calculated FRET efficiencies were determined for each of the ten independent 2 $\mu$s production simulations (Fig. S6). S2U2 exhibited a higher average FRET efficiency than S2U1 (0.846 $\pm$ 0.031 versus 0.805 $\pm$ 0.055, mean $\pm$ between-replicate standard deviation). In contrast, S2U1 exhibited greater within-replicate fluctuations in FRET efficiency, with an average standard deviation of 0.117 compared with 0.096 for S2U2. Bootstrap resampling across independent replicates yielded 95% confidence intervals that excluded zero for both the difference in mean FRET efficiency (0.004 to 0.078) and the difference in within-replicate standard deviation (-0.038 to -0.001). These findings indicate that although S2U1 and S2U2 were previously characterized as conformations within the same state-2 ensemble, they exhibit distinguishable FRET sampling behaviors across independent simulations.

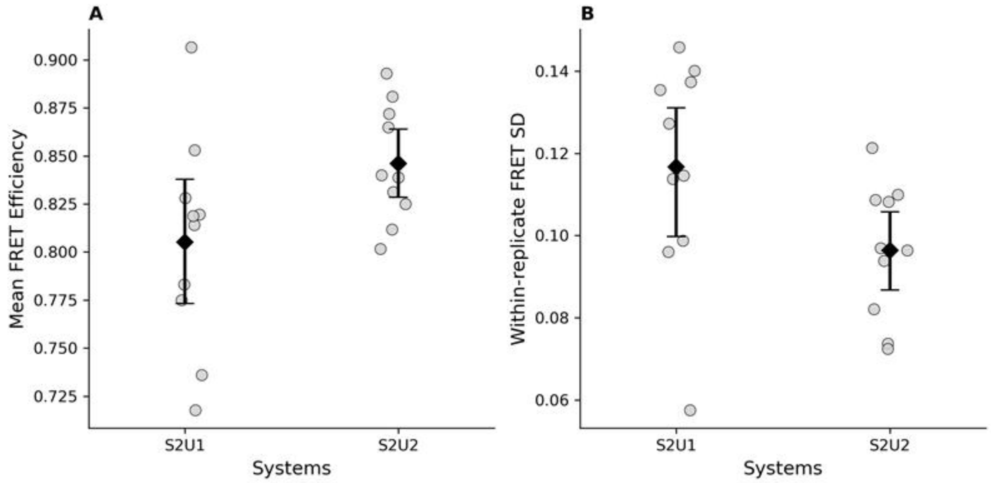


**Fig. S6.** Quantitative comparison of calculated FRET efficiencies across independent S2U1 and S2U2 simulations. (A) Mean FRET efficiencies and (B) within-replicate standard deviations of FRET efficiency calculated from ten independent 2 $\mu$s production simulations of each Env structure. Each gray circle represents an individual simulation, while black diamonds indicate the group means. Error bars represent 95% bootstrap confidence intervals obtained by resampling independent replicates over 10,000 iterations.

Additional PCA projections were analyzed to evaluate whether the conformational differences observed between S2U1 and S2U2 were maintained across multiple collective motion coordinates (Fig. S7). The first three principal components accounted for 52% of the total variance, with PC1, PC2, and PC3 contributing 26%, 14%, and 12%, respectively (Fig. S7A).

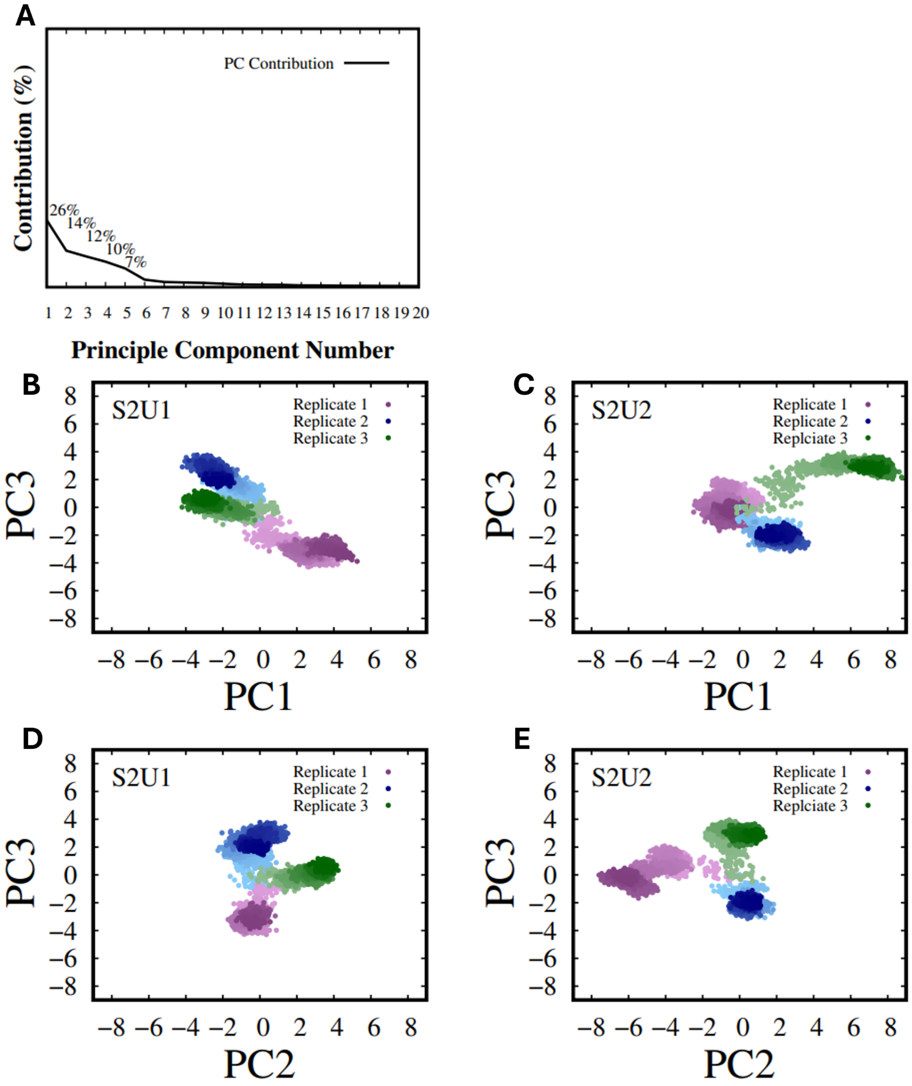


**Fig. S7.** Additional PCA projections of the extended uncleaved HIV-1 Env simulations. (A) Contribution of the first 20 principal components to the total variance, with the first three principal components accounting for 26%, 14%, and 12% of the total variance, respectively. (B,C) Projection of S2U1 and S2U2 trajectories onto PC1 and PC3. (D,E) Projection of S2U1 and S2U2 trajectories onto PC2 and PC3.

While the main text presents the projection along the two dominant modes (PC1 and PC2), additional projections involving PC3 were examined to characterize other major collective motions sampled during the simulations. The PC1-PC3 (Fig. S7B-C) and PC2-PC3 (Fig. S7D-E) projections demonstrate that S2U1 and S2U2 occupy distinct regions of principal component space across multiple collective motion coordinates. These results further support differences in the conformational ensembles sampled by the two Env states.

To characterize differences in trimer symmetry between the S2U1 and S2U2 conformational states, inter-protomer distances were monitored throughout the extended MD simulations (Fig. S8). Distances between corresponding gp120 and gp41 regions of neighboring protomers were calculated for each of the three independent replicas.

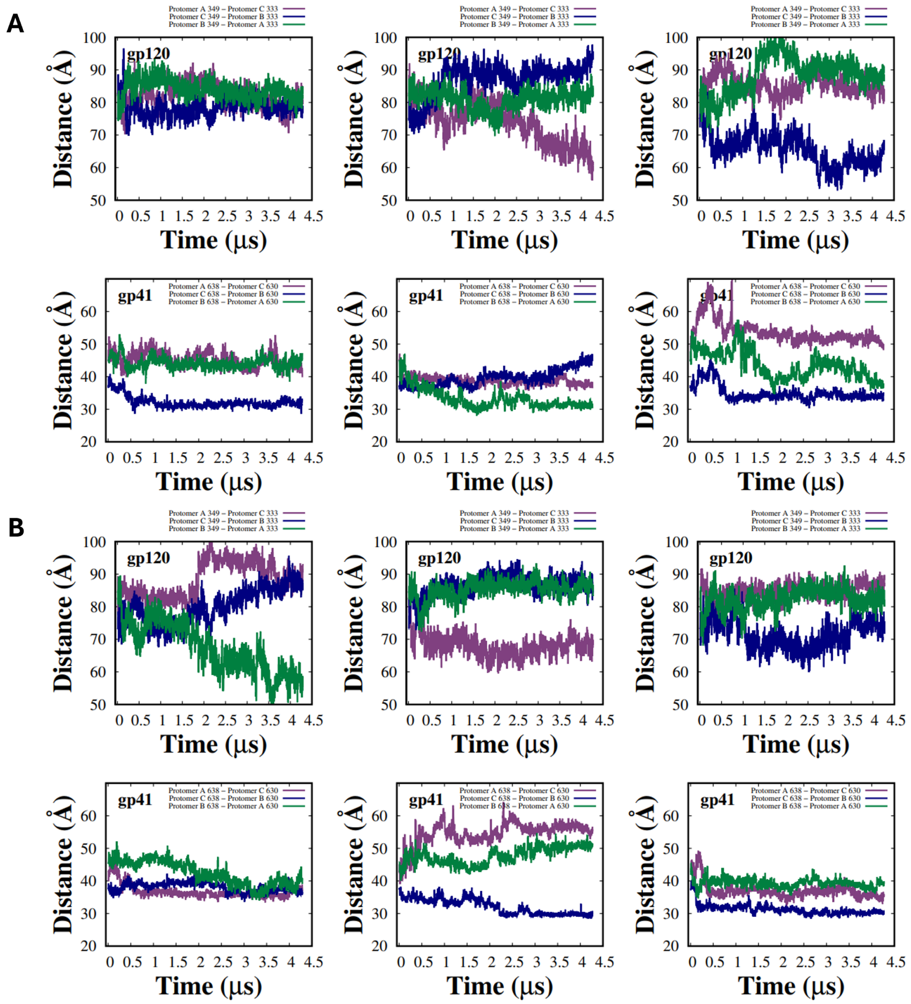


**Fig. S8.** Time evolution of inter-protomer distances used to quantify structural asymmetry in the extended uncleaved HIV-1 Env simulations. (A) S2U1 and (B) S2U2 distance trajectories from three independent ∼4.3 $\mu$s simulations. For each system, the upper row shows gp120 protomer-center distances and the lower row shows gp41 protomer-center distances. Columns correspond to independent replicas (left: Replicate 1, middle: Replicate 2, right: Replicate 3).

These distance measurements were subsequently used to calculate protomer asymmetry values, which quantify the deviation from an equivalent threefold symmetric arrangement.

To further characterize differences in quaternary organization between S2U1 and S2U2, the mean and standard deviation of gp120 and gp41 protomer asymmetry scores were calculated for each of the ten independent 2 $\mu$s production simulations (Fig. S9). S2U1 exhibited an average gp120 asymmetry score of 7.36 ± 2.76 Å (mean ± between-replicate standard deviation), compared with 7.86 ± 3.10 Å for S2U2. The corresponding gp41 asymmetry scores were 4.74 ± 3.01 Å and 4.80 ± 2.22 Å for S2U1 and S2U2, respectively. Bootstrap resampling across independent replicates yielded 95% confidence intervals that included zero for the differences in mean gp120 asymmetry (-1.91 to 2.96 Å) and gp41 asymmetry (-2.15 to 2.25 Å). Similarly, the differences in within-replicate asymmetry fluctuations were not clearly resolved between the two structures. These findings indicate that the average protomer asymmetry and its fluctuations were broadly comparable between S2U1 and S2U2 during the initial 2 $\mu$s simulations.

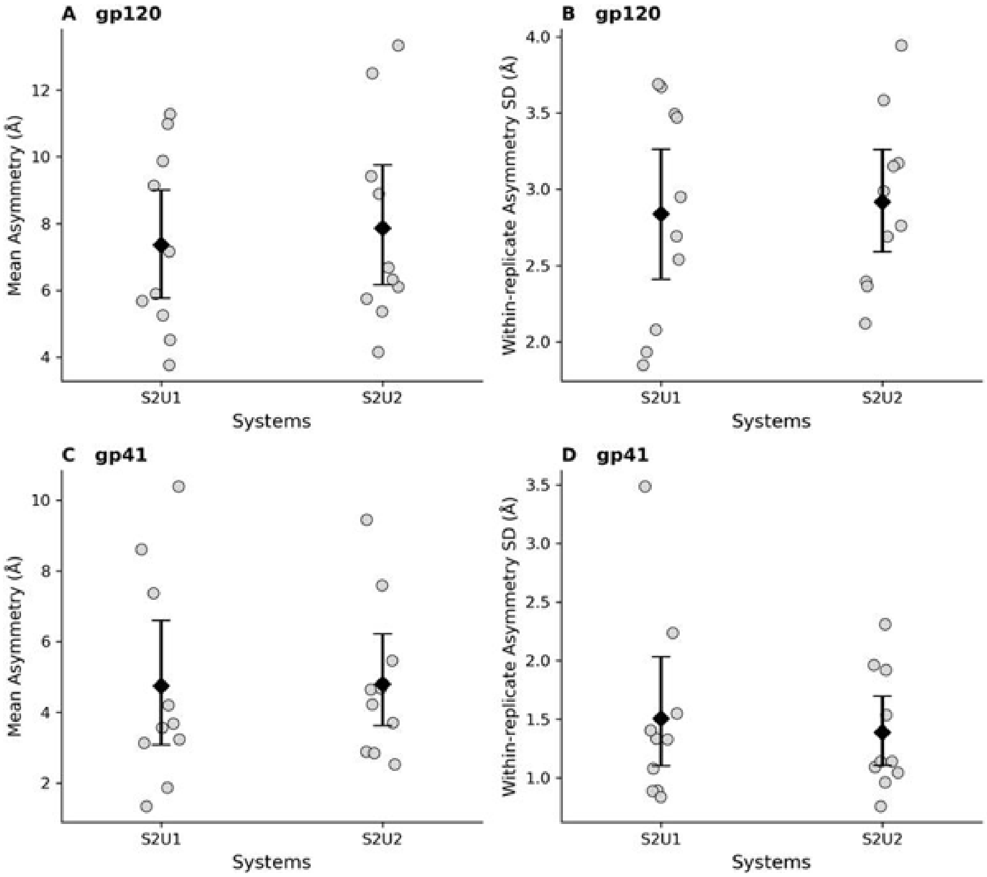


**Fig. S9.** Quantitative comparison of gp120 and gp41 protomer asymmetry across independent S2U1 and S2U2 simulations. Mean protomer asymmetry scores for gp120 (A) and gp41 (C), and corresponding within-replicate standard deviations for gp120 (B) and gp41 (D), calculated from ten independent 2 $\mu$s production simulations of each Env structure. Each gray circle represents an individual simulation, while black diamonds indicate the group means. Error bars represent 95% bootstrap confidence intervals obtained by resampling independent replicates over 10,000 iterations. Asymmetry scores were calculated as the sample standard deviation of three equivalent inter-protomer distances at each trajectory frame.
